# Building molecular band-pass filters via molecular sequestration

**DOI:** 10.1101/2022.04.02.486709

**Authors:** Yichi Zhang, Christian Cuba Samaniego, Katelyn Carleton, Yili Qian, Giulia Giordano, Elisa Franco

## Abstract

Engineered genetic circuits with tailored functions that mimic how cells process information in changing environments (e.g. cell fate decision, chemotaxis, immune response) have great applications in biomedicine and synthetic biology. Although there is a lot of progress toward the design of gene circuits yielding desired steady states (e.g. logic-based networks), building synthetic circuits for dynamic signal processing (e.g. filters, frequency modulation, and controllers) is still challenging. Here, we provide a model-based approach to build gene networks that can operate as band-pass filters by taking advantage of molecular sequestration. By suitably approximating the dynamics of molecular sequestration, we analyze an Incoherent Feed-Forward Loop (IFFL) and a Negative Feedback (NF) circuit and illustrate how they can achieve band-pass filter behavior. Computational analysis shows that a circuit that incorporates both IFFL and NF motifs improves the filter performance. Our approach facilitates the design of sequestration-based filters, and may support the synthesis of molecular controllers with desired specifications.

## I. Introduction

Beyond sensing and responding to molecular and physical inputs, cells have the ability to detect and react to *temporal* changes in said inputs [1], [2]. Temporal patterns are essential in routine processes such as cell differentiation, death, and even controlling the immune response. To process temporal signals at the molecular level, it has been found that cells take advantage of recurring motifs, like Negative Feedback (NF), Positive Feedback (PF), and Incoherent Feed-Forward Loops (IFFL) [3], [4]. Because this organization bears many similarities to classical designs in electronic signal-processing circuits, an exciting research direction is the composition of these motifs to systematically build artificial genetic networks for processing dynamical signals.

IFFL motifs have been used in previous works as core architectures for the design of fold change detectors [5], pulse generators [6], gradient sensing mechanisms [7], and to achieve disturbance rejection [8]. In parallel, NF motifs have guided the design of controllers [9], oscillators [10], and pulse generators [11]. These demonstrations suggest that the combination of IFFL and NF motifs may enable the synthesis of arbitrary, more complex dynamic circuits and signal processors.

In this paper, through theoretical analysis and numerical computations, we evaluate the signal processing capabilities of combined IFFL and NF motifs. Using these motifs, we suggest routes to build molecular filters that take direct inspiration from well-known architectures in analog filter design. In section II-A and II-B, we first focus on the IFFL and NF motif in isolation to operate as a band-pass filter. Further, in section II-C, we combine both motifs to overcome design challenges for band-pass filter. Finally, a brief discussion on filter design in section III.

## II. Results

In previous works, we demonstrated how an IFFL motif with delay or a negative feedback network can operates as a practical derivative operator [7], [12]. Here, we first study both network’s capacity to operate as band-pass filter independently. Later, we analyze a network that combines both motifs (IFFL and NF) to address the limitation of their single motifs.

### A. An IFFL-based network operates as a band-pass filter

An incoherent feed-forward loop processes a single input *U* and up-regulates outputs protein *Y* and mRNA *Z* at rate constants *k* and *β*, respectively. Then, *Z* produces protein *X* at a rate constant *θ*. In addition, *X* down-regulates *Y* through a sequestration mechanism at a rate constant *γ/ξ*. The parameter *ξ* is a non-dimensional parameter that tunes the sequestration rate. Finally, species *X, Y, C* and *Z* decay at rate constants *δ* and *ϕ*. Fig. 1A shows the the IFFL circuit, associated with the chemical reactions:

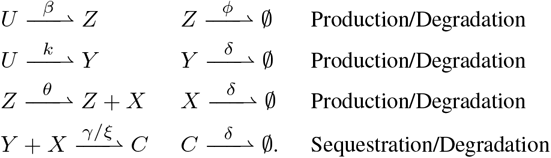

Using the law of mass action, we can write the Ordinary Differential Equations (ODEs) describing the dynamics of species concentrations:

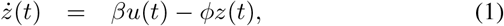

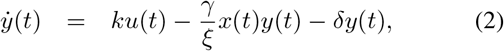

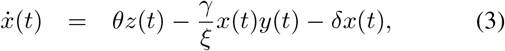

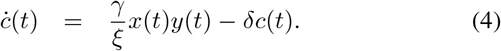

We can show that (1)-(4) is a positive system.

#### Proposition 1

Since *u*(*t*) *>* 0 for all *t*, system (1)-(4) is positive: *z*(0), *y*(0), *x*(0), *c*(0) ≥ 0 implies *z*(*t*), *y*(*t*), *x*(*t*), *c*(*t*) ≥ 0 for all *t >* 0.

*Proof:* When *z*(*t*) = 0, then *ż*(*t*) ≥ 0, and the same holds for *y*(*t*), *x*(*t*) and *c*(*t*), which proves positivity. ▪

**Fig. 1.**
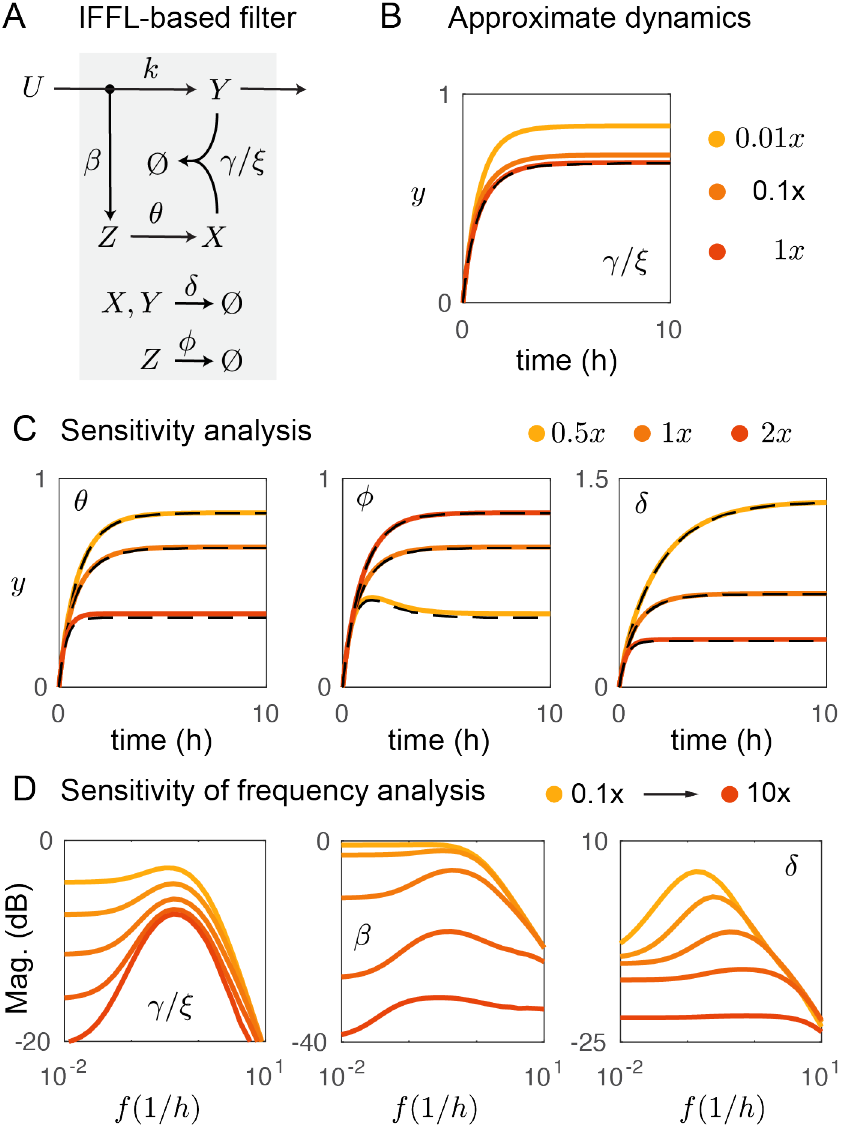
IFFL-based band-pass filter. A) Circuit schematic based on molecular sequestration. B) The full model’s dynamics converge to the approximate dynamics, dash black line. C) Comparing prediction of approximate dynamics for different parameter rates. Nominal parameters (NP): *β* = *k* = *θ* = *δ* = 1, *ϕ* = 3, and *γ/ξ* = 100. D) Frequency response as a function of network’s parameter. The same as before except for *θ* = 1*/*5, and *ϕ* = *βθ/k*.

#### 1) Approximate dynamics in the fast sequestration regime

Equations (1)-(4) include nonlinear terms. To overcome the challenges due to nonlinearity, we introduce the new states *x*^*T*^ (*t*) = *x*(*t*) + *c*(*t*) and *y*^*T*^ (*t*) = *y*(*t*) + *c*(*t*), representing the total amount of the respective species *X* and *Y* that is either free or bound to the complex *C*. Then, we can rewrite equations (2)-(4) equivalently as

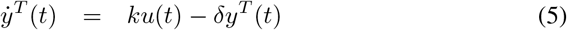

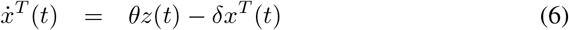

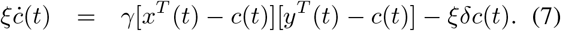

The trajectories of system (1), (5)-(7) are bounded within an invariant set, which contains at least an equilibrium point.

##### Proposition 2

The set

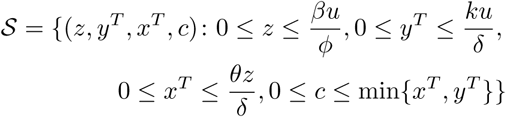

is positively invariant for the system (1), (5)-(7), namely, (*z*(*t*_0_), *y*^*T*^ (*t*_0_), *x*^*T*^ (*t*_0_), *c*(*t*_0_)) ∈ 𝒮 implies (*z*(*t*), *y*^*T*^ (*t*), *x*^*T*^ (*t*), *c*(*t*)) ∈ 𝒮 for all *t* ≥ *t*_0_. Therefore, the system admits an equilibrium point in 𝒮.

*Proof:* When *y*^*T*^ (*t*) = 0, then 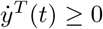, and the same holds for *x*^*T*^ (*t*), *c*(*t*) and *z*(*t*), thus guaranteeing the lower bounds. As for the upper bounds, when 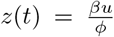, then *ż*(*t*) ≤ 0; when 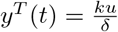, then 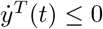; when 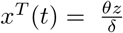, then 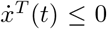. When *c*(*t*) = min {*x*^*T*^ (*t*), *y*^*T*^ (*t*)}, then we can have three cases: if *x*^*T*^ (*t*) *< y*^*T*^ (*t*), then invariance is guaranteed by Nagumo’s theorem provided that 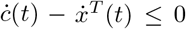, which is true because 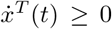 within 𝒮; if *x*^*T*^ (*t*) *> y*^*T*^ (*t*), then it must be 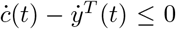, which is again true because 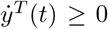 within 𝒮; if *x*^*T*^ (*t*) = *y*^*T*^ (*t*), then both 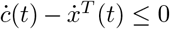 and 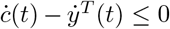 must hold, which is indeed the case. Positive invariance of the compact set 𝒮 implies that 𝒮 includes an equilibrium point [13]. ▪ Allowing *c*(*t*) *>* min {*x*^*T*^ (*t*), *y*^*T*^ (*t*)} would mean that either *x*(*t*) or *y*(*t*), or both, are negative, which is unacceptable since they represent concentrations.

Then we can consider the equilibrium value 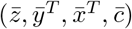, where 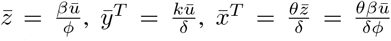 and 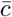 can be computed as a solution to 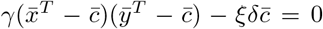. In a fast sequestration regime, *ξ* → 0. Since the roots of a polynomial are continuous functions of the coefficients, the limit of 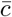 for *ξ* → 0 is the *smallest* solution to the algebraic equation 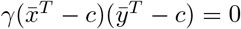. Hence,

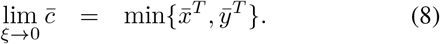

Then, the limit of the asymptotic value of *y* = *y*^*T*^ − *c* in the fast sequestration regime can be computed as

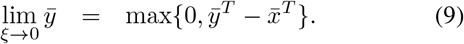

It can also be shown that, for *ξ* sufficiently small, the differences *e*_*c*_(*t*) = | min*{x*^*T*^ (*t*), *y*^*T*^ (*t*)*}* − *c*(*t*)| and *e*_*y*_(*t*) = | max*{*0, *y*^*T*^ (*t*) − *x*^*T*^ (*t*)*}* − *y*(*t*)| are upper bounded for all *t* ≥ *T*, with *T* appropriately chosen depending on the initial conditions: as *t* increases, the differences asymptotically decrease to 0. The smaller *ξ >* 0, the slower the dynamics of *x*^*T*^ and *y*^*T*^ with respect to the dynamics of *c*; hence, *x*^*T*^ and *y*^*T*^ can be seen as approximately constant in the equation of *ċ*. The faster the dynamics of *c*(*t*) (i.e., the smaller *ξ >* 0), the faster the transient after which the differences *e*_*c*_ and *e*_*y*_ become negligible. This legitimates the use of max {0, *y*^*T*^ (*t*) − *x*^*T*^ (*t*)} to approximate *y*(*t*).

Since the dynamics (1) of *z*, (5) of *y*^*T*^ and (6) of *x*^*T*^ are described by linear ODEs, we can derive their transfer functions in the Laplace domain and write:

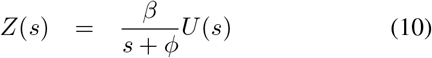

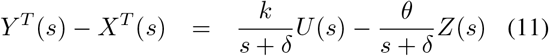

By substituting *Z*(*s*) from (10) into (11), we get

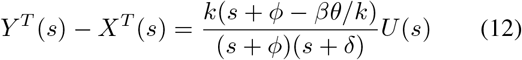

The transfer function in (12) does not describe the whole dynamics of the full system, but it can be used to compute the approximation of *y*(*t*) as

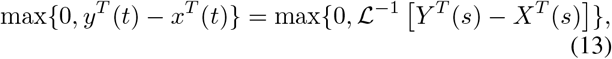

where ℒ^−1^ denotes the inverse Laplace transform.

In Fig. 1B, we compare the time evolution of *y*(*t*) for different sequestration rates *γ/ξ* (increasing from yellow to red), by solving equations (1)-(3) for a step input *u* (starting from zero initial conditions), and also its approximation (dash black line) obtained by evaluating (13). The trajectories of the full model quickly converge to expression (13) for larger *γ/ξ* (the two curves are very close at all times because the transient is negligible, being the dynamics of *c*(*t*) very fast and the initial conditions zero). We also compare the evolution of *y*(*t*) obtained by solving the full system for different rates *θ, ϕ*, and *δ*, along with its approximation, as shown in Fig. 1C. The simulations confirm that, in the fast sequestration regime, we can successfully approximate the dynamics of *y*(*t*) by evaluating expression (13).

The discrepancy *e*_*y*_ between approximated and full-dynamics trajectories is larger when *y* takes smaller values. These discrepancies decrease for even larger sequestration rates (not shown here).

#### 2) Frequency response in the fast sequestration regime

Transfer functions are useful to understand the frequency response for *linear* systems. Although the IFFL-based network is nonlinear due to the sequestration reaction, in the fast sequestration regime we can use (13) to get an approximation of the frequency response, and not only of the time evolution.

Fig. 1D shows the frequency analysis of the IFFL-based circuit by solving computationally equations (1)-(3) for a periodic input *u* = 0.5(*sin*(*ft*) + 1). When *ϕ* = *θβ/k*, the transfer function (12) has a zero at the origin (*s* = 0). This is a key feature of the IFFL-based network, which enables us to tune the zero of the system. Ideally, this results in a band-pass filter, as shown in the right side of Fig. 1D. We evaluate the frequency response for different values of sequestration rate. Smaller values of *γ/ξ* (yellow lines) result in the absence of sequestration (less effective subtraction part), which leads to a first order frequency response, while larger values of sequestration (red line) lead to a well-defined band-pass filter.

To further understand how the system parameters affect the frequency response, we vary the parameters *β* and *δ* as show in Fig. 1D. By either increasing or decreasing *β*, it will change the magnitude of the zero in the numerator. As a result, the frequency response will converge to a constant magnitude for low frequencies. The cut-off frequency experiences some small changes. On the other hand, *δ* changes the frequency cut-off of the band pass filter for small values. The effects of these parameters can also get to similar conclusion by analyzing the transfer function from equation (12).

#### 3) Frequency response of the linearised system

Consider the frequency response of the IFFL-based circuit linearised around its equilibrium and expressed in the shifted variables 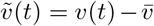, where *v* = (*z, y, x*) and 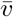 is the equilibrium state, and *ũ*(*t*) = *u*(*t*) − *ū*, where *ū* is the equilibrium input. The linearised system is therefore 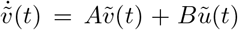, and the shifted output is 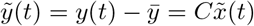, with

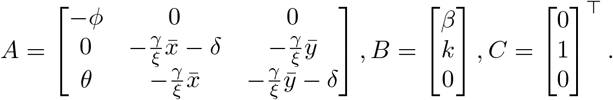

We can write the transfer function from *U* (*s*) to *Y* (*s*), *G*(*s*) = *C*(*sI* − *A*)^−1^*B*, as

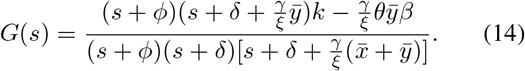

In the fast sequestration regime, considering a small *ξ* leads to the approximation

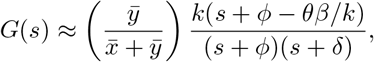

which matches the transfer function in (12) when 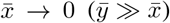, hence the first term becomes (approximately) one.

An IFFL motif can therefore be used to build a band-pass filter, provided that the parameters are carefully tuned in a fast sequestration regime and *ϕ* ≈ *θβ/k*. A critical aspect is that the cut-off frequencies of the band-pass filter depend on the degradation rates *ϕ* and *δ*, which could be challenging to tune in practice.

### B. Band-pass filter based on negative feedback (NF)

We propose a negative-feedback motif (NF-based circuit) that realizes a band-pass filter with tunable frequency as shown in Fig. 2A. An input *U* produces at rate constant *k* species *Y*, which at rate constant *ρ* produces *Z*, which in turn produces *X* at rate constant *θ*. Species *X* sequesters *Y* at rate constant *γ/ξ*, where the non-dimensional parameter *ξ* tunes the sequestration rate. Species *Z* decays at rate *ϕ*, while *Y, X*, and *C* all decay at rate *δ*. The corresponding chemical reactions are:

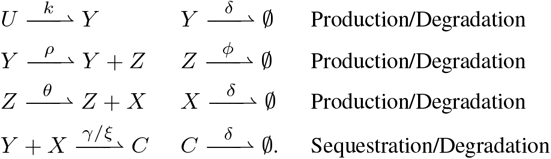

**Fig. 2.**
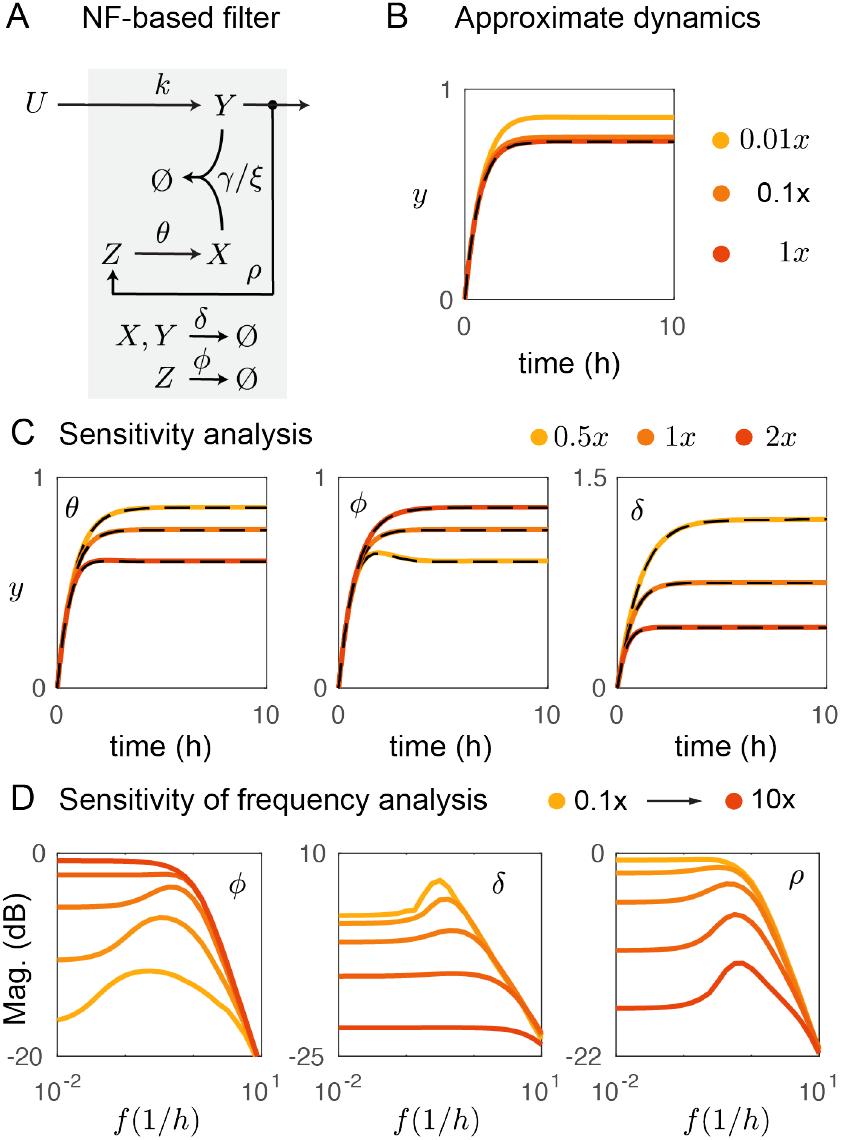
NF-based band-pass filter. A) Negative feedback circuit. B) Approximate dynamics as a function of *γ*. C) Comparing approximation dynamics (black dash line) with the full system. NP: *ρ* = *k* = *θ* = *δ* = 1, *ϕ* = 3, and *γ/ξ* = 100. D) Computing the frequency response for periodic inputs. The same as before except for *θ* = 1*/*5, and *ϕ* = *βθ/k*.

We use the law of mass action to derive the ODE dynamical model:

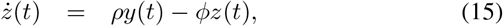

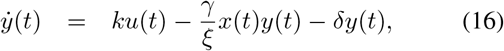

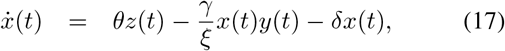

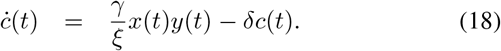

#### 1) Approximate dynamics in the fast sequestration regime

Introducing the new state variables *x*^*T*^ (*t*) = *x*(*t*) + *c*(*t*) and *y*^*T*^ (*t*) = *y*(*t*) + *c*(*t*) leads to the ODEs:

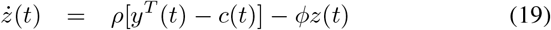

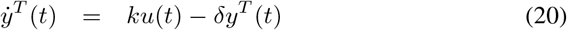

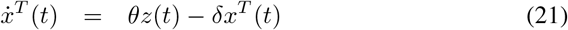

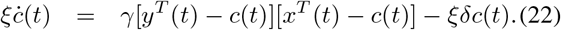

Following the same approach as in the previous section, we can prove that system (15)-(18) is positive and the set

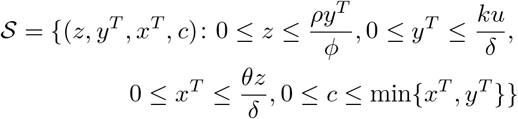

is positively invariant for system (19)-(22), which therefore admits an equilibrium point in 𝒮. We cannot allow *c*(*t*) *>* min {*x*^*T*^ (*t*), *y*^*T*^ (*t*)}, because either *x*(*t*) or *y*(*t*), or both, would become negative.

At the equilibrium, 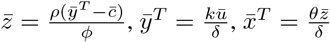 and 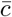 can be computed as a solution to 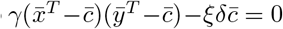. Therefore, in a fast sequestration regime, *ξ* → 0,

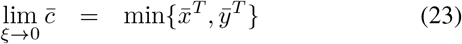

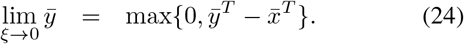

For *ξ* sufficiently small, we can make the same considerations as in the previous section about the differences *e*_*c*_(*t*) and *e*_*y*_(*t*): the smaller *ξ >* 0, the faster the dynamics of *c*(*t*), which allows us to neglect the transient. After the (very fast) transient, the differences *e*_*c*_ and *e*_*y*_ become negligible, which legitimates the use of max {0, *y*^*T*^ (*t*) − *x*^*T*^ (*t*)} as an approximation of *y*(*t*).

Hence, for *large enough times t*, the dynamics of *y*^*T*^ − *x*^*T*^ can be approximated by

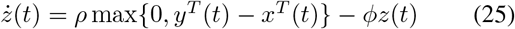

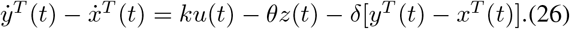

Considering the approximate equations under the additional assumption that *y*^*T*^ ≥ *x*^*T*^ and moving to the Laplace domain leads to the transfer function

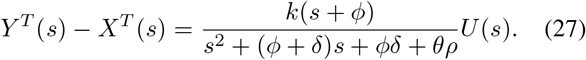

In Fig. 2B, considering a step input *u*, we vary the sequestration rate and compare the evolution of *y*(*t*) computed by solving equations (15)-(17), shown in yellow to red lines, with its approximation from expression (27), shown in black dashed lines. As expected, when for the sequestration rate *γ/ξ* is large (i.e. *ξ* is small), *y*(*t*) is very close to its approximation. We compare exact and approximated trajectories for different parameters *ϕ, δ*, and *ρ*, as shown in Fig. 2C. In all our numerical tests, the approximation is very good in the fast sequestration regime: the larger the values of the sequestration rate, the smaller the discrepancy.

#### 2) Frequency response in the fast sequestration regime

Equation (27) can be regarded as the approximated transfer function for the NF-based circuit under fast sequestration, for positive values of *y*^*T*^ − *x*^*T*^. Examining the transfer function (27) suggests that the NF-based circuit can behave as an ideal band-pass filter if *ϕ* = 0. However, this requirement can be challenging to implement experimentally. Moreover, for small values of *ϕ*, the system can operate as a band-pass filter and also have a constant gain for low frequencies, as shown in Fig. 2D. Smaller values of *δ* and larger values of the feedback gain *ρ* shows improve the band-pass behavior.

#### 3) Frequency response of the linearised system

As done in the previous section, we now consider the frequency response of the NF-based circuit linearised around its equilibrium. The linearised system has matrices

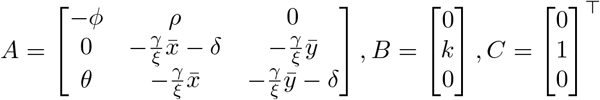

corresponding to the transfer function from *U* (*s*) to *Y* (*s*)

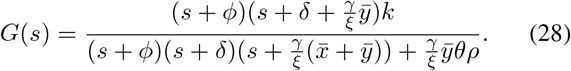

In the fast sequestration regime, assuming *ξ >* 0 very small and 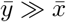 leads to the approximation

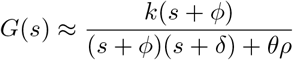

which matches the expression in equation (27). This suggests that, in the fast sequestration regime, the linearised system well approximates the dynamics of the original nonlinear system, also in terms of frequency behavior. With an appropriate choice of the parameter values, the NF-based circuit behaves as a band-pass filter whose cut-off frequencies can be tuned by adjusting the strength of the overall feedback. However, a drawback of this circuit is that it cannot deliver an ideal band-pass filter.

### C. Band-pass filter based on incoherent feed-forward loop (IFFL) and negative feedback (NF)

We now take advantage of both IFFL-based and NF-based designs to obtain a band-pass filter with tunable cut-off frequencies. Fig. 3A illustrates the IFFL/NF-based circuit, showing all the involved species and their interactions. An input *U* produces *Y* at rate *k* and *Z* at rate *β*. Species *Y* produces *Z* at rate *ρ*, while *Z* produces *X* at rate *θ*. Species *X* is sequestered by *Y* at rate *γ/ξ* (*ξ* is a non-dimensional parameter). Species *Z* decays at rate *ϕ*, while *Y, X*, and *C* all decay at rate *δ*. The corresponding chemical reactions

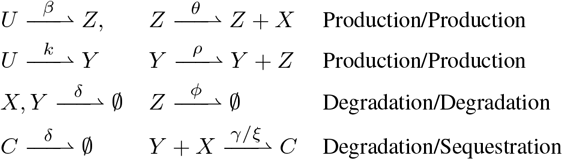

are associated, under the law of mass action, with the dynamical ODE model

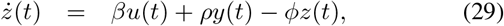

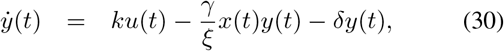

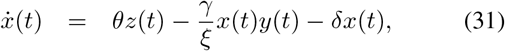

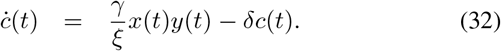

**Fig. 3.**
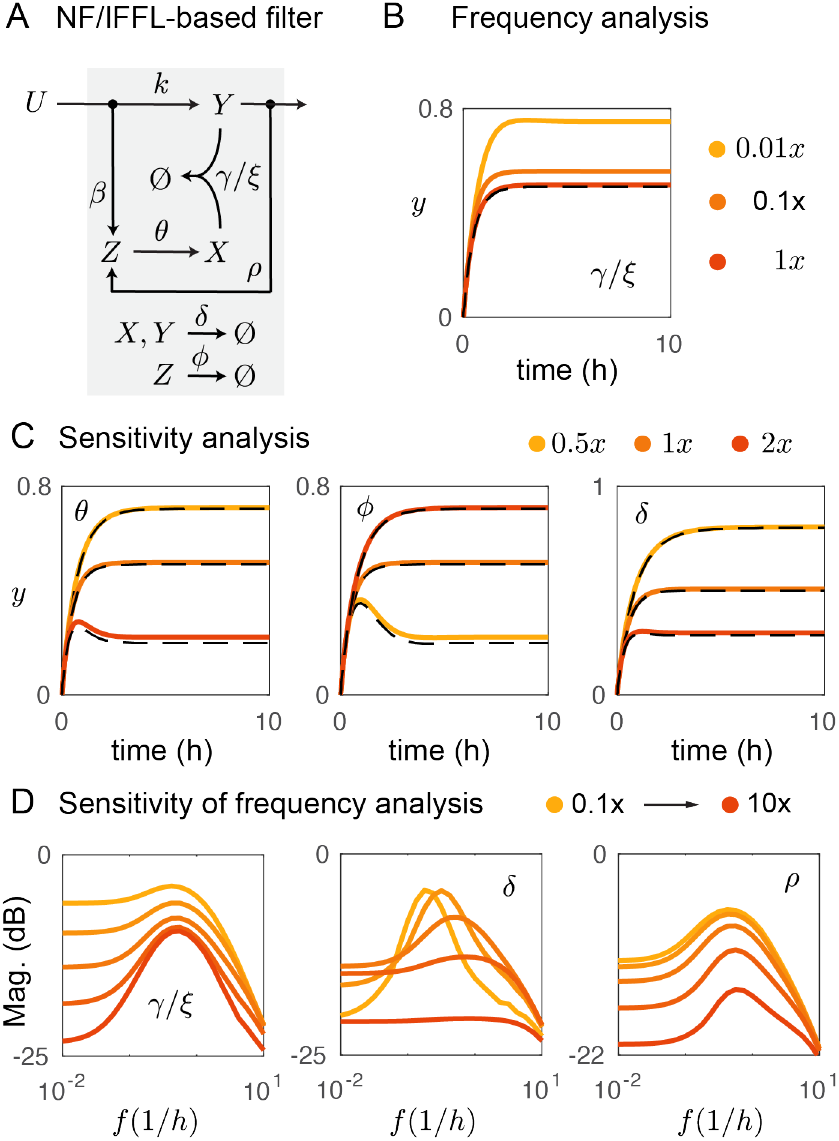
NF/IFFL-based band-pass filter. A) Circuit schematic. B) Approximation in the fast sequestration regime. C) Comparing prediction by varying different parameters. NP: *β* = *ρ* = *k* = *θ* = *δ* = 1, *ϕ* = 3, and *γ/ξ* = 100 D) Characterization of frequency response. The same as before except for *θ* = 1*/*5, and *ϕ* = *βθ/k*.

#### 1) Approximate dynamics in the fast sequestration regime

Defining the new states variables *x*^*T*^ (*t*) = *x*(*t*) + *c*(*t*) and *y*^*T*^ (*t*) = *y*(*t*) + *c*(*t*) results in the ODEs:

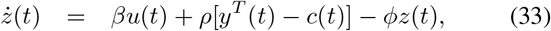

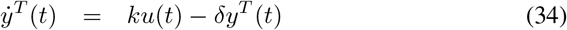

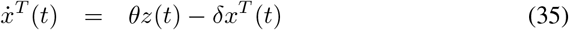

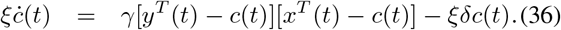

Following the same approach as in the previous sections, we can prove that system (29)-(32) is positive and the set

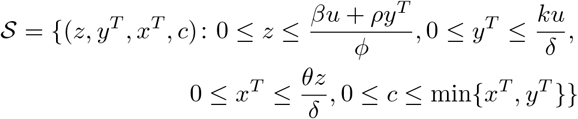

is positively invariant for system (33)-(36), which therefore admits an equilibrium point in 𝒮. As before, we need to rule out *c*(*t*) *>* min {*x*^*T*^ (*t*), *y*^*T*^ (*t*)} to prevent *x*(*t*) and *y*(*t*) from becoming negative.

At the equilibrium, 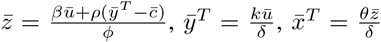 and 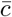 solves 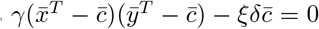. Therefore, in a fast sequestration regime, *ξ* → 0,

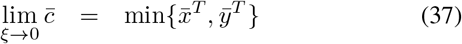

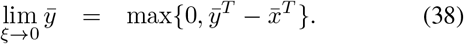

Again, the smaller *ξ >* 0, the faster the dynamics of *c*(*t*), which allows us to neglect the transient and approximate *y*(*t*) as max*{*0, *y*^*T*^ (*t*) − *x*^*T*^ (*t*)*}* for *t* large enough, when also the dynamics of *y*^*T*^ − *x*^*T*^ can be approximated by

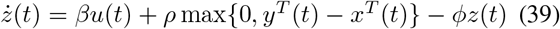

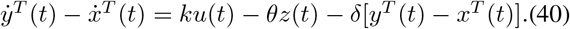

Considering the approximate equations under the additional assumption that *y*^*T*^ ≥ *x*^*T*^ and moving to the Laplace domain leads to the transfer function

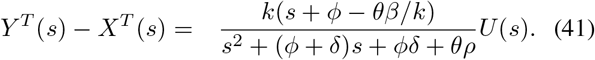

Fig. 3B compares the exact evolution of *y*(*t*) in the IFFL/NF-based circuit, computed from (29)-(31), for differ sequestration rates (from yellow to orange), with its approximation provided by expression (41). The discrepancy is small for large sequestration rates. We vary different parameters *θ, ϕ* and *δ* to compare the performance of the exact and approximate dynamics in Fig. 3C.

#### 2) Frequency response in the fast sequestration regime

We computationally characterize the frequency response of the IFFL/NF-based circuit for fast sequestration rates according to the approximated transfer function (41). An ideal band-pass filter admits a zero at the origin (*s* = 0), which can be obtained by setting *ϕ* = *θβ/k*. Fig. 3D shows the frequency response for different sequestration rates. A well defined band-pass filter behavior is obtained for large values of *γ/ξ* (red line). With small sequestration rates, we get a first order frequency response (yellow line). The band-pass behavior is improved for small *δ* and we can vary *ρ* to tune the frequency response.

#### 3) Frequency response of the linearised system

Again, we consider the frequency response of the system linearised around its equilibrium. The IFFL/NF-based circuit has linearised dynamics described by matrices

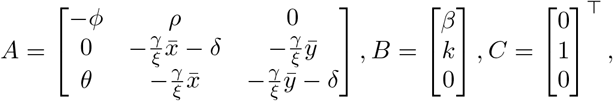

corresponding to the transfer function

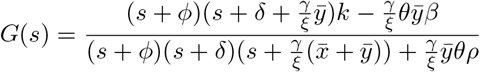

from *U* (*s*) to *Y* (*s*). In the fast sequestration regime, assuming *ξ >* 0 very small and 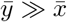 yields the approximation

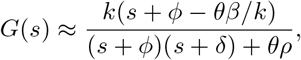

which matches the approximation in (41) and can guide the design of a band-pass filter with the desired transfer function.

The IFFL/NF-based circuit leverages the main advantages of both motifs (IFFL and NF). The IFFL motif allow us to achieve a zero at the origin (*s* = 0) by setting *ϕ* ≈ *θβ/k*, while the NF motif allows us to tune the cut-off frequencies by tuning the strength of the negative feedback *θρ*. Combining both features considerably improves the band-pass filter performance.

## III. Discussion

In this paper, we suggest a model-based approach to engineer molecular networks that exhibit band-pass filter behavior by taking advantage of molecular sequestration. Our findings on the design of synthetic genetic networks for dynamical molecular filters expand the repertoire of functions of IFFL (e.g fold change detection [5], pulse behavior [6], gradient sensing [7], and NF motifs (e.g feedback controllers [9], gradient sensing [12], oscillators [10], pulse generator [11]). It also expands and complements the experimental effort to engineer signal processing based on transcriptional regulation [14], [15], [16]. A limitation of the NF-based filter is that the sequestering species must have zero degradation rate constant. Such a requirement is challenging to meet in practical settings. The IFFL-based network implements a subtraction operation to deliver an ideal band-pass filter, however at low sequestration rate this operation is poorly approximated. In summary, both individual motifs present a non-ideal band-pass filter, but their combined tuning within a single circuit can improve the filter behavior.

The approach we introduce allows us to systematically study the dynamics of sequestration-based circuits in either open- or closed-loop systems. We note that the transfer functions of the IFFL and NF networks resemble a Proportional (P) and Derivative (D) controller. This indicates that our filters may facilitate the design of PD feedback controllers with a target frequency response. We also foresee that these circuits may help to realize biomolecular recurrent neural networks, building on previously demonstrated sequestration-based neural networks [17]. Because a key feature of sequestration-based networks is their ultrasensitive and tunable response [18], we speculate that ultrasensitivity may be key for building signal processors. For this reason, the synthesis of molecular filters may be possible with other ultrasensitive mechanisms distinct from sequestration.

